# Reduced subcortical envelope phase locking in middle age without detectable changes in perceptual discrimination

**DOI:** 10.64898/2026.11.20.768042

**Authors:** David López-Ramos, Victor Bauer, Laurel H. Carney, Emmanuel Ponsot

## Abstract

Cochlear synaptopathy (CS) can occur during normal aging without elevated audiometric thresholds and has been hypothesized to contribute to suprathreshold auditory difficulties. Frequency-following responses (FFRs) have been proposed as markers of CS, but whether their envelope- and temporal-fine-structure-related components are similarly affected by age, and whether such changes relate to perceptual sensitivity, remains unclear. Here, we examined associations between age-related FFR changes and perceptual sensitivity in young (n = 20, 19-32 years) and middle-aged (n = 20, 40-58 years) adults with clinically normal hearing. Participants completed psychoacoustic discrimination tasks for formant frequency, formant bandwidth, and temporal-envelope shape using stationary stimuli in quiet and background noise. FFRs were recorded using dynamic stimuli varying along the same acoustic dimensions. We quantified envelope- and temporal-fine-structure-related phase locking (PLV_ENV_ and PLV_TFS_) across harmonics and assessed how it tracked stimulus changes over time. Mean PLV_ENV_ decreased significantly with age, and PLV_ENV_ tracking of decreasing formant bandwidth and temporal-envelope shape weakened with age. In contrast, mean PLV_TFS_ and temporal slopes of PLV_TFS_ spectral centroid were not significantly associated with age. Discrimination thresholds showed no significant associations with age, either in quiet or in noise, and PLV measures did not significantly explain individual differences in discrimination thresholds. These findings indicate that age-related reductions in subcortical envelope phase locking can be detected by middle age without corresponding measurable reductions in perceptual sensitivity.

**Significance statement:** Cochlear synaptopathy (CS) is increasingly invoked as a contributor to age-related auditory decline, but its functional consequences in humans remain poorly understood. Using electrophysiological and psychophysical measures in young and middle-aged adults with clinically normal hearing, we found age-related reductions in subcortical phase locking to sound envelopes, whereas phase locking to temporal fine structure was not significantly related to age. Psychophysical discrimination thresholds, in quiet or in noise, were not significantly related to age or to any of the phase locking metrics. These findings suggest that early age-related reductions in envelope phase locking, a candidate marker of early CS, need not be accompanied by measurable impairments in basic auditory discrimination.

## 1 Introduction

Cochlear synaptopathy (CS), the loss of synapses between inner hair cells and auditory-nerve (AN) fibers, can occur without elevated audiometric thresholds (Kujawa and Liberman, 2009; Kohrman et al., 2020). Histological studies in humans suggest that 30-50% of AN fibers are lost between 20 and 60 years of age despite normal audiometric sensitivity (Wu et al., 2019), making middle age a window of substantial yet clinically hidden neural degeneration. This loss may impair suprathreshold processing (Bramhall et al., 2019, 2024; Oxenham, 2016) and contribute to speech-in-noise difficulties (Garrett et al., 2025). Proposed CS biomarkers include auditory brainstem responses, middle-ear muscle reflexes and frequency-following responses (FFRs) (Liberman et al., 2016; Vasilkov et al., 2021; Märcher-Rørsted et al., 2022; Bharadwaj et al., 2022), but their relationship to synaptic loss in humans remains uncertain, complicating estimates of its prevalence and impact (Bramhall et al., 2019; Bharadwaj et al., 2019; Liu et al., 2024; Kampel et al., 2026; Heassler et al., 2026).

The FFR is a promising marker. The FFR reflects AN and brainstem activity (Bidelman, 2018; Krizman and Kraus, 2019) evoked by sustained complex sounds, offering a bridge to perception, and its envelope (FFR_ENV_) and temporal fine structure (FFR_TFS_) components reflect neural phase locking to these two aspects of sound, respectively (Aiken and Picton, 2008). FFR_ENV_ reductions have been reported with aging in animals (Parthasarathy and Kujawa, 2018; Zink et al., 2025), after experimentally induced CS (Shaheen et al., 2015; Garrett et al., 2025), and in humans with suspected CS, including middle-aged adults (Ponsot et al., 2025; Zink et al., 2025) and older listeners (Vasilkov et al., 2021; Garrett et al., 2025). FFR_TFS_ to tones and vowel-like sounds also appears to decline with age, with evidence from studies including middle-aged and older adults (Märcher-Rørsted et al., 2022, 2026; Ponsot et al., 2025). Whether FFR_ENV_ and FFR_TFS_ are similarly affected by middle age remains unclear. FFR-based CS markers typically sum response magnitudes across harmonics into a single scalar metric, using stationary stimuli (Vasilkov et al., 2021; Ponsot et al., 2025), but combining FFR across harmonics may obscure differences in their sensitivity to acoustic conditions, perceptual performance and aging. For example, noise has opposite effects on the first two harmonics of FFR_ENV_ (Billings et al., 2020), only harmonics 3-5 combined predict speech-in-noise performance (Mepani et al., 2021), and age-related FFR_TFS_ reductions were driven by the harmonic nearest the spectral-envelope peak (Ponsot et al., 2025). It remains unclear whether FFRs at different harmonics are affected differently by aging in middle-aged adults, and a detailed characterization of these age-related changes is needed.

The perceptual consequences of age-related CS remain uncertain. Age-related FFR reductions have been observed without psychoacoustic deficits (Ponsot et al., 2025), whereas age-related perceptual deficits have been reported without consistent changes in FFR_ENV_ (Patro et al., 2021). Associations between CS markers and speech-in-noise performance remain inconsistent (DiNino et al., 2022; Bramhall and McMillan, 2024), despite positive findings (Garrett et al., 2025; Zink et al., 2025). Clarifying these inconsistencies may require assessing psychoacoustic performance under challenging listening conditions, such as in background noise (DiNino et al., 2022), and matching stimuli across FFR and psychoacoustic measures in the same listeners (Garrett et al., 2025).

Here, we investigated whether age-related FFR changes relate to perceptual performance in young and middle-aged normal-hearing adults. We used dynamic stimuli varying in formant frequency, formant bandwidth, and temporal-envelope shape to characterize early age effects in envelope-and temporal-fine-structure-related phase locking (PLV_ENV_ and PLV_TFS_) across harmonics. The same stimuli were used to measure psychoacoustic discrimination thresholds in quiet and background noise. We hypothesized that aging would affect both ENV and TFS coding, accompanied by reduced perceptual sensitivity. Instead, we found significant age-related reductions in PLV_ENV_ but not in PLV_TFS_ measures. Psychoacoustic performance was not affected by age, and FFR measures did not significantly explain individual differences in discrimination thresholds.

## 2 Methods

### 2.1 Participants, session protocol and audiological assessment

#### 2.1.1 Participants and session protocol

Fifty-three participants were recruited through the Institut Européen d’Administration des Affaires (INSEAD) on the basis of age and self-reported normal hearing. Recruitment targeted young adults aged 18-35 years and middle-aged adults aged 40-60 years. Eligibility required audiometric thresholds ≤25 dB HL at all frequencies from 0.125 to 8 kHz in the test ear and at least 90% correct responses in each of the three psychoacoustic screening tasks. Otoscopy was performed to rule out visible abnormalities of the outer ear and obstruction of the external auditory canal.

Twelve participants (1 young and 11 middle-aged adults) were excluded because they did not meet the audiometric criterion, and one middle-aged participant withdrew from the study. The final sample comprised 40 participants: 20 young normal-hearing adults (yNH; 10 males; mean age ± SD, 23.1 ± 4.0 years) and 20 middle-aged normal-hearing adults (mNH; 10 males; 49.3 ± 5.4 years). All participants in the final sample successfully completed the psychoacoustic screening tasks. Participants also completed a non-standardized questionnaire on neurological conditions, self-reported auditory disorders, and noise exposure. Responses to this questionnaire were not included in the statistical analyses.

The core experimental protocol comprised four sessions conducted over three or four days, totaling approximately 8.5 h per participant (Sessions 1 -4 in **Fig. 1A**). The initial screening session lasted approximately 1 h and included otoscopy, pure-tone audiometry, and training of the psychoacoustic discrimination tasks. The main experiment comprised three subsequent sessions of approximately 2.5 h each. During each session, participants first completed adaptive psychoacoustic measurements of formant-frequency, formant-bandwidth, and envelope-shape discrimination thresholds, followed by electrophysiological recordings. All psychoacoustic and electrophysiological measurements during these three sessions were conducted in the same sound-attenuated booth, with stimuli presented monaurally to the test ear selected during the audiological assessment. When two sessions were conducted on the same day, they were separated by at least 3 h.

**Figure 1.**
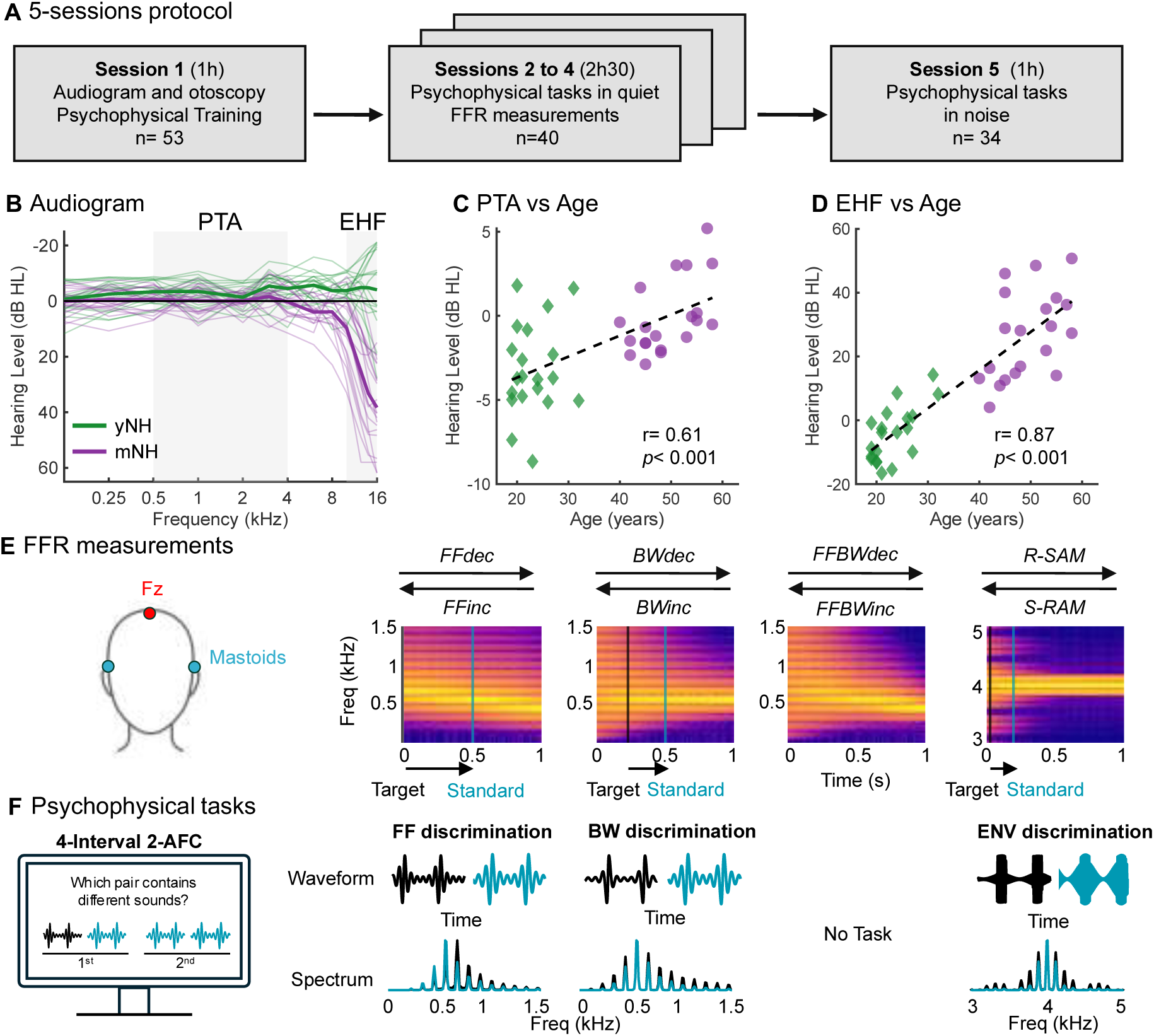
Session protocol, audiometric characteristics of the participants, and overview of electrophysiological and psychoacoustic measurements. ***A***, Experimental timeline and participant flow. Session 1 comprised otoscopy, pure-tone audiometry, and psychoacoustic screening/training (n = 53). Participants meeting the inclusion criteria completed three main experimental sessions (Sessions 2-4; n = 40), each including psychoacoustic measurements in quiet and FFR recordings. A subset of participants completed an additional session involving the psychoacoustic tasks in noise (Session 5; n = 34). ***B***, Individual pure-tone audiograms for young normal-hearing (yNH; green) and middle-aged normal-hearing adults (mNH; purple) from conventional through extended high frequencies; thick lines indicate group averages. ***C***, ***D***, Relationship between age and pure-tone average (PTA; mean hearing threshold at 0.5, 1, 2, and 4 kHz; ***C***) and extended-high-frequency threshold (EHF; mean threshold at 10, 12.5, 14, and 16 kHz; ***D***). Dashed lines indicate linear fits; correlation coefficients and associated *p* values are shown in each panel. ***E***, Electrode configuration and dynamic stimuli used for FFR measurements. The analysis signal was obtained from Fz referenced to the mean of the two mastoid electrodes. Spectrograms illustrate dynamic changes in formant frequency (FF), formant bandwidth (BW), combined FF and BW, and envelope shape for the high-frequency amplitude-modulated stimuli. Arrows indicate the two opposite stimulus trajectories: *FFdec*/*FFinc*, *BWdec*/*BWinc*, combined *FFBWdec*/*FFBWinc*, and rectangular-to-sinusoidal/sinusoidal-to-rectangular amplitude modulation (*R-SAM*/*S-RAM*). Target and standard configurations used in the corresponding psychoacoustic tasks are indicated schematically. ***F***, Psychoacoustic discrimination paradigm and stimuli. Thresholds were measured using a four-interval, two-alternative forced-choice paired-comparison task in which participants identified the pair containing two different sounds. Example waveforms and spectra illustrate the target (black) and standard (cyan) stimuli for FF, BW, and envelope-shape (ENV) discrimination.

A subset of 34 participants (14 yNH and 20 mNH) completed an additional 1-h session involving psychoacoustic discrimination tasks in noise (Session 5 in **Fig. 1A**). During this additional session, each participant completed the psychoacoustic tasks alone in a separate quiet, isolated booth.

The study was approved by the INSEAD Ethics Committee and conducted in accordance with the Declaration of Helsinki. Participants were informed about the experimental procedures, provided written informed consent before participation, and received financial compensation.

#### 2.1.2 Audiological assessment

Air-conduction thresholds were measured bilaterally from 0.125 to 16 kHz using the iAudiogram system and RadioEar DD450 circumaural headphones, which were calibrated before data collection. Measurements were conducted in a sound-attenuated booth. Participants were instructed to press a response button whenever they detected a sound. Thresholds were estimated using a Bayesian adaptive procedure (Schlittenlacher et al., 2018), with a targeted estimated precision of ±2.45 dB. These measurements were obtained using a newly automated pure-tone audiometry application developed by My Medical Assistant SAS (iAudiogram, www.iaudiogram.com). This algorithm is inspired by recent machine-learning based audiometry (Schlittenlacher et al., 2018), has been validated against state-of-the-art audiologist-based manual audiometry, and has already been used for research purposes (Jean et al., 2025; Wallaert et al., 2025).

The procedure began with an initialization phase in which conventional audiometric frequencies from 0.125 to 12 kHz were tested over a maximum of 50 trials. This was followed by a Bayesian adaptive phase comprising 50-84 trials, in which frequency and presentation level were selected to minimize uncertainty in the threshold estimate. Thresholds were represented as continuous functions of frequency for each ear (**Fig. 1B**). Extended-high-frequency thresholds above 8 kHz were retained for subsequent analyses but were not used to determine participants’ eligibility. The ear with the lower pure-tone average (PTA) was selected as the test ear for all subsequent psychoacoustic and electrophysiological measurements. PTA was calculated across 0.5, 1, 2, and 4 kHz, whereas the extended-high-frequency threshold (EHF) was calculated across 10, 12.5, 14, and 16 kHz. Both PTA and EHF measures were significantly higher in mNH compared to yNH (PTA: Welch’s t_(36.32)_= −3.94, *p*< 0.001; EHF: Welch’s t_(31.85)_= −8.58, *p*< 0.001); these measures were also significantly positively associated with age (**Fig. 1C,D**)

### 2.2 Electrophysiological measurements

#### 2.2.1 Stimuli

Two stimulus families were used: single-formant harmonic complexes adapted from Allen and Oxenham (2014) and amplitude-modulated tones, targeting low-frequency and high-frequency cochlear regions, respectively. The former allowed independent manipulation of formant frequency and formant bandwidth, whereas the latter varied in envelope shape along a continuum between rectangular amplitude modulation (RAM) and sinusoidal amplitude modulation (SAM). Dynamic stimuli were used for electrophysiological (EEG) measurements (**Fig. 1E**), whereas stationary stimuli defined along the same acoustic dimensions were used for psychoacoustic measurements (**Fig. 1F**).

The single-formant stimuli were spectrally shaped harmonic complexes with a fundamental frequency (F0) of 113 Hz. The spectral envelope was defined by two parameters: formant frequency and formant bandwidth, the latter expressed as the spectral-envelope slope in dB/octave. More negative slopes produced narrower spectral peaks. Six dynamic trajectories were created from these stimuli, comprising three pairs with opposite temporal directions (**Fig. 1E**). For the formant-frequency trajectories, the bandwidth remained fixed at -12 dB/octave. Formant frequency decreased linearly from 678 to 452 Hz in *FFdec* and increased linearly from 452 to 678 Hz in *FFinc*. For the bandwidth trajectories, formant frequency remained fixed at 565 Hz, so that it corresponded exactly to the fifth harmonic. The bandwidth changed linearly from -3 to -21 dB/octave in *BWdec* and from -21 to -3 dB/octave in *BWinc*. Two additional trajectories varied formant frequency and bandwidth simultaneously. In *FFBWdec*, formant frequency decreased linearly from 678 to 452 Hz while the bandwidth changed from -3 to -21 dB/octave. The reverse trajectory, *FFBWinc*, increased formant frequency from 452 to 678 Hz while the bandwidth changed from -21 to -3 dB/octave.

Two additional dynamic trajectories were generated using amplitude-modulated tones with a 4-kHz carrier and a modulation frequency of 113 Hz, matching the F0 of the single-formant stimuli (**Fig. 1E**). Envelope shape was defined by a blending coefficient, m:

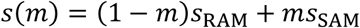

where s_RAM_ and s_SAM_ denote the RAM and SAM tones, respectively; m = 0 corresponds to RAM and m = 1 to SAM. In the RAM-to-SAM trajectory (*R-SAM*), the coefficient changed over time according to

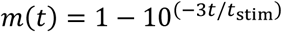

where t_stim_ is the 1-s stimulus duration. This exponential trajectory was selected to produce an approximately linear change, in decibels, in the amplitudes of the stimulus harmonic components. The reverse trajectory (*S-RAM*; **Fig. 1E**) traversed the same continuum in the opposite temporal direction.

All electrophysiological stimuli were generated with custom-made MATLAB software (R2024b, The MathWorks, Inc) at a sampling rate of 44.1 kHz, lasted 1 s, and were gated with 12.5-ms onset and offset Tukey ramps. Root-mean-square amplitude was normalized within each F0 period for the single-formant stimuli and within each modulation period for the amplitude-modulated stimuli to maintain stimulus energy over time.

#### 2.2.2 EEG recording and procedure

Electrophysiological responses were recorded using a BioSemi ActiveTwo system (BioSemi, Amsterdam, The Netherlands). Stimuli were delivered through an RME Fireface UCX II audio interface at a sampling rate of 44.1 kHz and 16-bit resolution and were presented monaurally through ER-2 insert earphones (Etymotic Research). The presentation system was calibrated at 1 kHz using the ER-2 earphone coupled through a 2-cm³ coupler to a Brüel & Kjær Artificial Ear Type 4153 connected to a Brüel & Kjær Model 2250 sound level meter. A second output from the audio interface was routed directly to the BioSemi system to record the presented stimulus and enable temporal alignment between stimulus onset and the EEG recordings.

Five electrodes were used: flat electrodes placed over the left and right mastoids (Lmon and Rmon), a pin-type electrode at Fz, and the Common Mode Sense (CMS) and Driven Right Leg (DRL) electrodes. The skin over the mastoids was cleaned with alcohol, and conductive gel was applied to ensure adequate electrode contact. The signal used for analysis was obtained from Fz referenced to the mean of the two mastoid electrodes (**Fig. 1E**).

Each of the eight dynamic stimuli was presented at 62.5 dB SPL for 840 repetitions, with each repetition comprising one positive-and one negative-polarity presentation. Opposite polarities were used to separate envelope-and temporal-fine-structure-related response components (Aiken and Picton, 2008). The 840 repetitions were distributed across six blocks, with two blocks completed in each of the three electrophysiological sessions. Each block contained 140 repetitions of each stimulus, and the order of the eight stimulus trajectories was randomized across blocks.

Each stimulus presentation lasted one second and was followed by a fixed 120-ms interval plus a random jitter drawn from a uniform distribution between 0 and 80 ms. Positive-and negative-polarity presentations alternated throughout the recording. A 10-s interval separated successive stimulus trajectories, and participants were allowed to rest between blocks.

#### 2.2.3 EEG preprocessing

EEG preprocessing and electrophysiological analyses were performed offline. After referencing Fz to the mean of the two mastoid signals, stimulus onsets were identified from the simultaneously recorded stimulus channel. The EEG signal was then segmented into time-aligned epochs separately for positive- and negative-polarity presentations.

Artifact rejection was based on the peak-to-peak amplitude of each epoch. An epoch was rejected when its peak-to-peak amplitude differed from the local median by more than three median absolute deviations, with the local distribution estimated from the 200 closest neighboring epochs. On average, 74 ± 37 epochs were rejected per trajectory and participant, corresponding to approximately 8.9% of the recorded epochs and leaving an average of 743 repetitions per trajectory. Retained epochs were high-pass filtered at 80 Hz using a 1500^th^-order finite impulse response (FIR) filter and low-pass filtered at 3 kHz using a 240^th^-order FIR filter.

#### 2.2.4 Frequency-following response analysis

In the present study, we preferred to assess phase-locking in the frequency-following responses using information available from the phase-locking value (PLV) spectrum (Zhu et al., 2013; Bharadwaj et al., 2014) rather than the magnitude (FFR) spectrum, because the former measure was less corrupted by noise. However, the two types of measures are expected to be strongly related (Zhu et al., 2013), and we verified that similar results were obtained if we had considered FFR_ENV_ and FFR_TFS_ instead of PLV_ENV_ and PLV_TFS_.

The bootstrap-based spectral analysis described by Zhu et al. (2013) was adapted to the dynamic stimuli by replacing the Fourier transform of each epoch with a short-time Fourier transform (STFT). STFTs were computed using 100-ms Blackman-Harris windows with 90% overlap and a 4096-point Fourier transform. For each bootstrap estimate, 400 epochs were randomly sampled. PLVs were estimated from 100 bootstrap draws. PLV quantified the consistency of response phase across repeated stimulus presentations. Values ranged from 0 to 1, with values approaching 1 indicating highly consistent phase across repetitions and values approaching 0 indicating little or no phase consistency (Zhu et al., 2013; Bharadwaj et al., 2014). Responses to opposite stimulus polarities were summed to obtain PLV_ENV_ and subtracted to obtain PLV_TFS_ (**Fig. 2A**).

**Figure 2.**
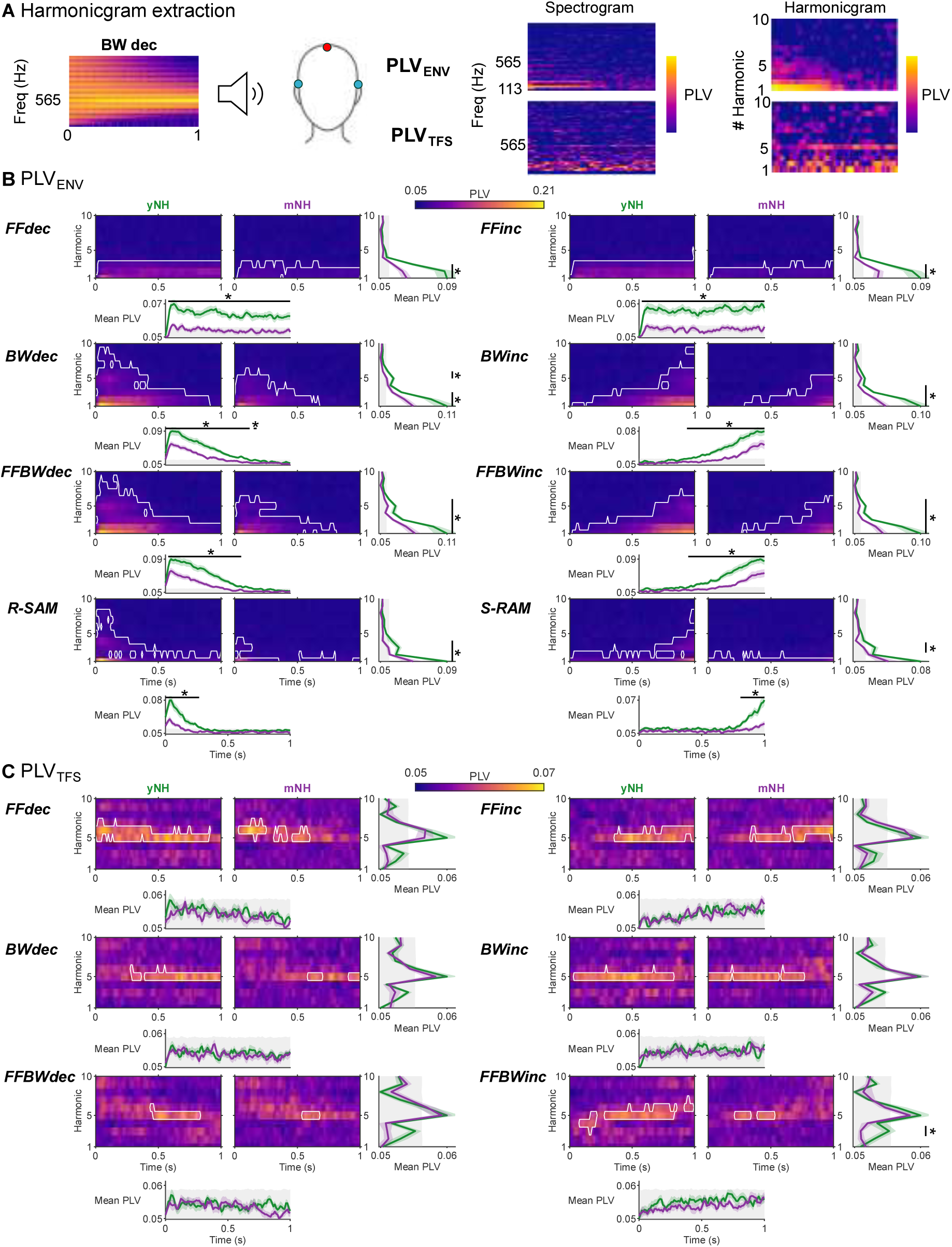
Envelope-and temporal fine-structure-related phase locking across dynamic stimulus trajectories. ***A***, Schematic overview of the extraction of harmonic-by-time PLV representations (harmonicgrams), illustrated for the decreasing-bandwidth (*BWdec*) stimulus in a representative participant. Envelope-and temporal fine-structure-related phase locking (PLV_ENV_ and PLV_TFS_) was quantified to obtain time-frequency PLV representations, from which PLV values at integer harmonics (1-10) of 113 Hz were extracted to construct the corresponding harmonicgrams. See ***Section 2.2.4*** for details of PLV estimation, and harmonicgram extraction. ***B***, Envelope-related phase locking (PLV_ENV_) for young normal-hearing (yNH) and middle-aged normal-hearing adults (mNH) across the eight stimulus trajectories: decreasing and increasing formant frequency (*FFdec*, *FFinc*), bandwidth (*BWdec*, *BWinc*), combined formant frequency and bandwidth (*FFBWdec*, *FFBWinc*), and the RAM-to-SAM and SAM-to-RAM transitions (*R-SAM*, *S-RAM*). For each trajectory, adjacent heatmaps show group-mean PLV across harmonics 1-10 over the 1 s stimulus interval. White contours delineate clusters in which PLV significantly exceeded the corresponding group-specific simulated noise distribution (see ***Section 2.4.1***, for details). Harmonic marginals to the right show PLV averaged across time for each harmonic, whereas temporal marginals below show PLV averaged across harmonics 1-10 at each time bin. Green and purple curves indicate the yNH and mNH group means, respectively, with shaded regions denoting ±1 SEM. Gray shading indicates the mean of the two groups’ 95^th^-percentile noise criteria. Black bars and asterisks indicate portions of significant between-group clusters that additionally met the display criterion that PLV in at least one group exceeded its own 95^th^-percentile noise criterion. ***C***, Temporal fine-structure-related phase locking (PLV_TFS_) for the six formant trajectories, displayed using the same conventions as in ***B***. Color denotes PLV magnitude, with a common color scale across trajectories within each panel and separate scales for PLV_ENV_ and PLV_TFS_.

Time-frequency PLV representations were converted into harmonicgrams by extracting PLV at integer multiples (from 1 to 10) of the fundamental frequency (Parida et al., 2021) (**Fig. 2A**). When a harmonic frequency did not coincide exactly with an STFT frequency bin, the larger PLV from the two adjacent bins was retained.

#### 2.2.5 PLV metrics

The harmonicgrams show PLV patterns across harmonics and time. We defined various scalar metrics to capture and test whether the overall trends that could be observed on the average harmonicgrams were statistically robust across listeners: mean PLV measures overall phase-locking strength, whereas temporal slopes describe changes in the electrophysiological response over the stimulus time course. Specifically, the temporal marginal slope summarizes changes in PLV_ENV_ over time, and the PLV_TFS_ spectral centroid slope summarizes changes in the distribution of PLV_TFS_ across harmonics over time.

### Mean PLV

For each participant and stimulus trajectory, mean PLV was calculated as the arithmetic mean of all harmonic-by-time values in the individual harmonicgram. The calculation included all available harmonics (1-10) and time bins, without restricting the calculation to significant clusters or values above the noise floor (see **Appendix**). Mean PLV_ENV_ was calculated for all eight trajectories. Mean PLV_TFS_ was calculated for the six trajectories involving changes in formant frequency, formant bandwidth, or both.

### Dynamic PLV_ENV_: Slope of the temporal marginal

Temporal changes in PLV_ENV_ were quantified from the participant-level harmonicgrams for all eight stimulus trajectories. At each time bin, PLV_ENV_ was averaged across harmonics H1-H10, to obtain a temporal marginal.

For each participant, the temporal marginal was smoothed with a 10-bin moving average, and its derivative was then calculated. Slopes are expressed in PLV/s, with positive and negative values indicating increasing and decreasing PLV_ENV_, respectively.

For each participant, a mean slope was calculated as the arithmetic mean of the signed derivative samples within 100-ms windows. For all the trajectories, we specifically examined the slope at the reference instant, which indicates the time at which each dynamic stimulus reached the acoustic value of the stationary, standard stimulus used in the corresponding psychoacoustic task. So, 100-ms analysis windows were positioned immediately before or after this time point, depending on trajectory direction, such that the change in the relevant acoustic dimension within the window matched the direction in which the target stimulus approached the standard stimulus during adaptive tracking in the psychoacoustic tasks (see **Section 2.3**). For *FFdec*, *BWdec*, and combined *FFBWdec*, the window was 0.4-0.5 s; for the corresponding increasing trajectories, it was 0.5-0.6 s. The reference instant was 0.5 s for these six trajectories. For *R-SAM*, the reference instant was approximately 0.2 s and the window was approximately 0.1-0.2 s; for *S-RAM*, the reference instant was approximately 0.8 s and the window was approximately 0.8-0.9 s. Thus, the window ended at the reference instant for decreasing trajectories and *R-SAM* and began at the reference instant for increasing trajectories and *S-RAM*.

### Dynamic PLV_TFS_: Slope of the spectral centroid

The PLV_TFS_ spectral centroid summarizes the distribution of PLV_TFS_ across harmonics at each time bin. For each participant and each of the six FF, BW, and combined FF and BW trajectories, the centroid was calculated from harmonics H1-H10 as:

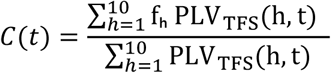

Here, fₕ = 113h Hz, spanning 113-1130 Hz. PLV_TFS_ values were used directly as weights, without noise-floor subtraction or significance-based selection. The resulting centroid, *C(t)*, is expressed in Hz and describes the weighted distribution of PLV_TFS_ across the fixed harmonic frequencies.

Each participant’s centroid trajectory was smoothed with a 10-bin moving average. A linear model, C(t) = a + bt, was then fitted by ordinary least squares across the full 1-s interval. The slope *b* was expressed in Hz/s and the intercept *a* represented the fitted centroid at t = 0.

Individual fit quality was screened separately for each trajectory using the root-mean-square error (RMSE) distribution pooled across both age groups. Fits with RMSE greater than Q3 + 1.5 IQR, where IQR=Q3-Q1, were excluded from participant-level age analyses and between-group comparisons. This screening removed three observations for *FFdec*, one each for *BWdec*, *BWinc*, and combined *FFBWinc*, and none for *FFinc* or combined *FFBWdec*. The resulting sample sizes were 37, 40, 39, 39, 40, and 39, respectively. Poor-fit participants were kept in the descriptive group centroid curves, which included all 20 participants per age group.

### 2.3 Psychoacoustic tasks

#### 2.3.1 Stimuli and paradigm

Discrimination thresholds in quiet were measured for formant frequency, formant bandwidth, and envelope shape, corresponding to the acoustic dimensions manipulated in the dynamic stimuli (**Fig. 1F**). Stationary stimuli used the same definitions as the stimuli used for EEG recordings, with parameters held constant over time. Psychoacoustic stimuli were delivered using the same calibrated audio interface and ER-2 insert-earphone system described for the electrophysiological recordings. All stimuli lasted 300 ms, had 30-ms onset and offset Tukey ramps, and were presented monaurally to the test ear at 62.5 dB SPL.

Each trial used a paired two-alternative forced-choice paradigm (**Fig. 1F**). Two stimulus pairs were presented sequentially: one contained two identical standard stimuli, and the other contained a standard and a target stimulus. Stimuli within a pair were separated by 200 ms of silence, and the pairs were separated by 600 ms. Participants selected the pair containing different sounds and received feedback after each trial. A similar paired-comparison paradigm has been used to measure formant-frequency discrimination (Carney et al., 2023).

#### 2.3.2 Formant-frequency discrimination

The reference was a single-formant harmonic complex with F0 = 113 Hz, formant frequency of 565 Hz, and spectral-envelope slope of -12 dB/octave. Only formant frequency differed between the standard and target; all other parameters were held constant. The initial target frequency was 790 Hz. This task assessed discrimination along the dimension varied in the dynamic FF trajectories used in EEG recordings.

#### 2.3.3 Formant-bandwidth discrimination

The single-formant stimuli had F0 = 113 Hz and a fixed formant frequency of 565 Hz. The standard bandwidth was -12 dB/octave, and the initial target bandwidth was -7.2 dB/octave. Only this bandwidth varied during adaptive tracking, assessing the dimension manipulated in the dynamic BW trajectories used in EEG recordings.

#### 2.3.4 Envelope-shape discrimination

Stationary hybrid RAM/SAM tones had a 4-kHz carrier and 113-Hz modulation frequency. The standard blending coefficient was m = 0.75, and the initial target coefficient was m = 0.21. During adaptive tracking, the comparison stimulus varied along the same RAM-SAM continuum used for the dynamic *R-SAM* and *S-RAM* stimuli used in EEG recordings.

#### 2.3.5 Screening, adaptive tracking, and run reliability

During the initial screening session, participants completed a 20-trial nonadaptive training run for each task. The target stimulus remained at the largest standard-target difference used to initiate the corresponding adaptive task. A score of at least 90% correct was required. Participants who did not meet this criterion could repeat the task, up to a total of four attempts; failure after four attempts resulted in exclusion.

During the three subsequent sessions, all tasks used a two-down, one-up adaptive procedure targeting approximately 70.7% correct performance (Levitt, 1971). The standard-target difference decreased after two consecutive correct responses and increased after each incorrect response. A common adaptive variable, X, expressed in decibels, was converted to the physical stimulus parameter using the following transformations.

Formant frequency:

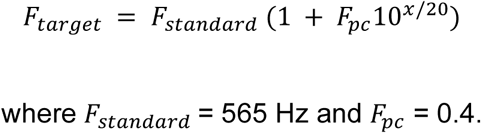

Formant bandwidth:

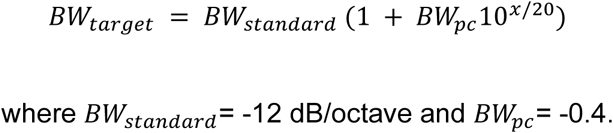

Envelope shape:

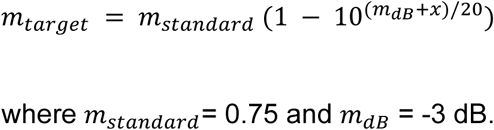

These transformations allowed the three physical stimulus parameters to be adjusted using a common decibel scale.

Step sizes were 6 dB for the first two reversals, 2 dB for reversals 3-6, and 1 dB for reversals 7-11. The threshold for each adaptive run was calculated as the mean of the final six reversal values. Catch trials comprised 10% of trials and presented the largest standard-target difference. They did not affect the adaptive track.

At least three adaptive runs were completed for each task in each session. An additional run was collected when the variability across the first three runs exceeded a predefined criterion. For formant-frequency and formant-bandwidth discrimination, an additional run was collected when the SD exceeded 30% of the mean threshold, consistent with Carney et al. (2023). For envelope-shape discrimination, an additional run was collected when the SD exceeded 6 dB.

FF thresholds are reported as percentage change in formant frequency relative to the reference value of 565 Hz. BW thresholds are expressed as percentage change relative to the reference value of 12 dB/octave. Envelope-shape discrimination thresholds are expressed in dB. Lower thresholds indicate better discrimination in all three tasks.

#### 2.3.6 Psychoacoustic task in noise

Psychoacoustic measurements in noise were conducted at the INSEAD-Sorbonne Université Behavioural Lab. Stimuli were generated using custom MATLAB software (R2024b, The MathWorks, Inc.) at a sampling rate of 48 kHz and 16-bit resolution and were presented monaurally to the same test ear used for the psychoacoustic measurements in quiet through Beyerdynamic DT 770 PRO headphones (250 Ω). Sound levels were calibrated using a Brüel & Kjær Model 2238 sound level meter and a custom-calibrated device consisting of a metal plate to which the headphones were coupled. The psychoacoustic stimuli were presented at 75 dB SPL together with pink noise band-limited from 100 Hz to 10 kHz. A new noise waveform was generated independently for each stimulus presentation. The simultaneous noise had the same 300-ms duration as the psychoacoustic stimulus and was gated with 30-ms onset and offset Tukey ramps. Signal-to-noise ratio (SNR) was defined from the RMS levels of the stimulus and noise and was set to 0 dB for the formant-frequency and formant-bandwidth discrimination tasks and +5 dB for the envelope-shape discrimination task. All other stimulus parameters, trial structure, and adaptive procedures were identical to those used for the corresponding psychoacoustic measurements in quiet.

Threshold in noise is reported in the same units as the threshold in quiet. The effect of noise is defined as the threshold in noise minus the threshold in quiet.

### 2.4 Experimental design and statistical analysis

The study used a mixed experimental design involving two age groups (yNH and mNH; n = 20 each). Electrophysiological measurements were obtained using eight dynamic stimulus trajectories, whereas psychoacoustic discrimination thresholds were measured for three acoustic dimensions in quiet (n = 40) and, in a subset of participants, in background noise (n = 34). Electrophysiological measures included mean PLV and metrics characterizing its temporal dynamics. Statistical analyses assessed differences between age groups, associations with age, and relationships between electrophysiological measures and psychoacoustic discrimination thresholds.

#### 2.4.1 Cluster-based assessment of PLV significance

To determine whether PLVs in each age group exceeded that expected under the noise model (see **Appendix**), group-mean PLV maps and marginal curves were assessed using one-sided cluster-mass tests adapted from Maris and Oostenveld (2007). For each group and stimulus trajectory, cells or bins exceeding the 95th percentile of their corresponding null distributions (200 realizations) were grouped into clusters using eight-neighbor connectivity for maps and consecutive bins for marginals. Cluster mass was calculated as the sum of the differences between observed PLV and mean null PLV across the cluster.

The same procedure was applied to each null realization, retaining only its maximum cluster mass to construct a null distribution of 200 maxima. The empirical *p* value of each observed cluster was calculated as the proportion of null maxima equal to or greater than its mass. Clusters with *p* < 0.05 were considered significant. Using the maximum cluster mass from each null realization provided correction for multiple comparisons across bins within each map or marginal.

#### 2.4.2 Cluster-based assessment of age-group differences in PLV

To determine whether PLV differed between yNH and mNH, harmonic and temporal marginals were compared using two-sided cluster-based permutation tests (Maris and Oostenveld, 2007), separately for each stimulus trajectory and PLV measure. At each bin, a t statistic was calculated, and consecutive bins exceeding an uncorrected threshold of *p*< 0.05 (df = 38) were grouped into separate positive and negative clusters. Cluster mass was calculated as the sum of the *t* statistics within each cluster.

To assess cluster significance, participants were randomly reassigned to the two age groups 5000 times, preserving group sizes (n = 20 each). For each permutation, the cluster analysis was repeated, and only the cluster with the largest absolute mass was retained, regardless of the direction of the group difference. The *p* value of each observed cluster was calculated as the proportion of permutations yielding a cluster with an equal or greater absolute mass. Clusters with *p*< 0.05 were considered significant. Using the largest cluster from each permutation provided correction for multiple comparisons across bins and both directions of the group difference within each marginal.

#### 2.4.3 Correlations and linear regression

Two-sided Pearson correlations were used to assess associations between age and the electrophysiological and psychoacoustic measures. For mean PLV, correlations were calculated by trajectory for PLV_ENV_ (all trajectories) and PLV_TFS_ (formant trajectories). Correlations between age and mean PLV_ENV_, mean PLV_TFS_, and the slope of the PLV_ENV_ temporal marginal were calculated using all 40 participants. Correlations between age and the PLV_TFS_ spectral-centroid slope included only participants who passed the trajectory-specific fit-quality screening (see **Section 2.2.5**).

For psychoacoustic performance, age correlations were calculated separately for FF, BW, and envelope-shape thresholds in quiet and in noise, and for the noise effect.

Correlation coefficients and two-sided *p* values were obtained using MATLAB’s corrcoef; the regression lines were calculated using polyfit with a first-degree polynomial.

#### 2.4.4 Linear mixed-effects models

Associations between PLV metrics and psychoacoustic thresholds were tested separately for FF, BW, and envelope-shape discrimination using linear mixed-effects models. The outcome was the threshold measured in quiet or noise, treated as a repeated observation; the noise-minus-quiet difference was not modelled as a separate outcome. Each model included a subject-specific random intercept.

Predictors were selected to match each task. For FF and BW, the models used mean PLV_ENV_, mean PLV_TFS_, the PLV_ENV_ temporal slope, and the PLV_TFS_ spectral-centroid slope, averaged across the corresponding decreasing and increasing trajectories. For envelope-shape discrimination, the predictors were mean PLV_ENV_ and the PLV_ENV_ temporal slope, averaged across *R-SAM* and *S-RAM*. For both slope measures, the predictor was the mean of the absolute values for the two trajectories.

Linear mixed-effects models were fitted by maximum likelihood separately for each task and PLV predictor, with discrimination threshold as the outcome. The contribution of listening condition (Quiet vs. Noise) was assessed by comparing an intercept-only model with a condition-only model. The contribution of age beyond condition was tested once per task by comparing the condition-only model with a model that also included age. The contribution of each PLV predictor beyond condition was assessed by comparing the condition-only model with a model that also included that predictor. The predictor’s contribution after adjustment for both condition and age was assessed by comparing the condition-plus-age model with a model that also included the PLV predictor. These nested models were compared using likelihood-ratio tests. Changes in Nakagawa marginal R² between the larger and smaller models are expressed as percentages of the total variance (100 × ΔR²). Linear mixed-effects models were fitted in R 4.6.1 using lme4 package (version 2.0-6; Bates et al., 2015). Marginal R² was calculated with performance package (version 0.18.2; Lüdecke et al., 2021), and base R’s stats functions were used for likelihood-ratio tests and p-value adjustment.

The primary analyses included 40 participants per task and up to 74 observations per task (40 quiet and 34 noise).

#### 2.4.5 Multiple-comparison correction

The Benjamini-Hochberg procedure was used to control the false discovery rate (FDR) (Benjamini and Hochberg, 1995), with significance set at an adjusted *p* value of < 0.05. For mean PLV, correction was applied separately to the eight age correlations across trajectories for PLV_ENV_ and the six correlations for PLV_TFS_. Age correlations with the PLV_ENV_ temporal slope and the PLV_TFS_ centroid slope were also corrected separately across their trajectory-specific tests, comprising eight and six tests, respectively.

For psychoacoustic age correlations, three families were defined by task (FF, BW, and envelope shape), each comprising the quiet threshold, noise threshold, and noise-minus-quiet effect. For the mixed-effects analyses, the condition contrasts formed one family of three tests, one for each task (FF, BW, and ENV). EEG-predictor comparisons were corrected separately for each task and model comparison: across four predictors for FF and BW and two predictors for ENV, both for the EEG contribution beyond condition and for the EEG contribution after adjustment for condition and age. The age-only model comparisons were not adjusted.

#### 2.4.6 Outlier detection and handling

Potential outliers were identified using Tukey’s 1.5 × IQR rule: values below Q1 -1.5 × IQR or above Q3 + 1.5 × IQR were flagged, where Q1 and Q3 are the 25th and 75th percentiles and IQR = Q3 -Q1. Outliers observed are shown as open symbols but they were retained in correlations, regression fits, FDR analyses, group summaries, and mixed-effects models.

## 3 Results

### 3.1 Age-related differences in neural phase locking to envelope and temporal fine structure across harmonics and time

The dynamic stimuli designed for this study allowed us to examine how PLV_ENV_ and PLV_TFS_ varied with stimulus features, and whether these phase-locking patterns differed between yNH and mNH listeners. **Figure 2B,C** shows group-mean PLV across harmonics 1-10 and throughout the one-second stimulus interval, together with harmonic and temporal marginal averages. PLV_ENV_ was assessed for all eight stimulus trajectories, whereas PLV_TFS_ was assessed for the six formant trajectories.

Significant clusters of PLV_ENV_ above the group-specific noise floor were observed for all eight stimulus trajectories in both age groups (cluster-corrected *p*< 0.05; **Fig. 2B**). In the harmonicgrams, responses to the *FFdec* and *FFinc* trajectories were concentrated predominantly within the lowest three harmonics throughout the stimulus interval in both groups. For *BWdec* and combined *FFBWdec*, significant clusters extended to higher harmonics early in the stimulus and became progressively restricted to lower harmonics towards the end, whereas the corresponding increasing trajectories showed the opposite pattern. Thus, the decreasing and increasing trajectories showed complementary patterns, with significant PLV_ENV_ extending to high harmonics during portions of the stimulus characterized by broader formant bandwidths. For *R-SAM*, significant PLV_ENV_ extended to high harmonics early in the stimulus, whereas for *S-RAM*, significant PLV_ENV_ clusters were observed at lower harmonics, extending to higher harmonics toward the end of the stimulus only in yNH. Thus, in both trajectories, PLV_ENV_ extended to higher harmonics as the modulation became more rectangular, consistent with the richer harmonic content of rectangular modulation (see **Fig. 1**). Consistent with these patterns, harmonic marginals in yNH showed significant above-noise PLV_ENV_ across harmonics 1-3 for both FF trajectories and *S-RAM*, 1-4 for *R-SAM*, 1-5 for *BWinc*, and 1-6 for *BWdec* and both combined FF and BW trajectories. In mNH, significant harmonic clusters spanned harmonics 1-5 for *BWdec* and harmonics 1-3 for the remaining formant trajectories, with an additional cluster at harmonic 5 for *FFBWdec*. For *R-SAM* and *S-RAM*, significant PLV_ENV_ in mNH was restricted to harmonics 1-2. Temporal marginals further showed that PLV_ENV_ remained significantly above noise across nearly the entire stimulus for both FF trajectories in yNH, whereas no significant temporal clusters were observed for these trajectories in the mNH group. For the BW and combined FF-BW trajectories, significant above-noise PLV_ENV_ was observed when the formant bandwidth was broader, corresponding to the earlier portion of the decreasing trajectories and the later portion of the increasing trajectories. These significant responses persisted over a larger portion of the stimulus in yNH than in mNH. For the high-frequency AM stimuli, significant PLV_ENV_ was observed when the modulation was more rectangular, around stimulus onset for *R-SAM* and near stimulus offset for *S-RAM* in both groups.

Between-group comparisons of PLV_ENV_ revealed significant differences between yNH and mNH for all eight stimulus trajectories (**Fig. 2B**). Only statistically significant between-group clusters, for which the PLV_ENV_ exceeded the corresponding noise criterion in at least one group, are reported (see ***Section 2.4.2***). In the harmonic marginals, PLV_ENV_ was significantly higher in yNH than in mNH at harmonics 1-3 for *FFdec* and *FFinc* (both *p*< 0.001), at harmonics 1-3 and 5-6 for *BWdec* (*p*= 0.007 and *p*= 0.032, respectively), and at 1-4 for *BWinc* (*p*= 0.001). PLV_ENV_ was also higher in yNH at harmonics 1-6 for *FFBWdec* (*p*= 0.001), *FFBWinc* (*p*< 0.001), at harmonics 1-4 for *R-SAM* (*p*= 0.001), and at harmonic 3 for S-RAM (*p*= 0.002). In the temporal marginals, higher PLV_ENV_ in yNH was observed across almost the entire stimulus interval for both FF trajectories (both *p*< 0.001). For the BW and combined FF-BW trajectories, significant between-group differences occurred when the formant bandwidth was broader, corresponding to the early and middle portions of the decreasing trajectories and the later portion of the increasing trajectories (all *p*< 0.001). For the high-frequency AM stimuli, significant between-group differences occurred when the modulation was more rectangular, around stimulus onset for *R-SAM* (*p*= 0.004) and near stimulus offset for *S-RAM* (*p*= 0.007).

Regarding PLV_TFS_, significant clusters above noise floor were observed for all six formant trajectories in both age groups (cluster-corrected *p*< 0.05; **Fig. 2C**). In the harmonicgrams, significant PLV_TFS_ was concentrated primarily around harmonics 5-6, with some clusters extending to adjacent harmonics. This region corresponded to the spectral location of the formant peak, indicating that the strongest PLV_TFS_ was concentrated near the formant peak. For *FFdec*, significant PLV_TFS_ clusters spanned harmonics 5-7 across most of the stimulus interval in yNH, whereas mNH showed separate clusters during the early and middle portions of the stimulus. For *FFinc*, significant PLV_TFS_ was observed mainly during the middle and later portions of the stimulus, spanning harmonics 5-6 in both groups. For *BWdec*, significant clusters spanned harmonics 5-6 from the early-to-middle portion of the stimulus through stimulus offset in yNH, whereas in mNH two later clusters were restricted to harmonic 5; for *BWinc*, significant PLV_TFS_ clusters were concentrated around harmonic 5, with limited extension to harmonic 6, during the early and middle portions of the stimulus in both groups. For *FFBWdec*, significant PLV_TFS_ clusters were more temporally restricted and were observed mainly around harmonics 5-6 in yNH and at harmonic 5 in mNH during the middle portion of the stimulus. For *FFBWinc*, significant PLV_TFS_ in yNH was observed first around harmonics 3-5, later around harmonics 5-6, and near stimulus offset around harmonics 6-7, whereas in mNH two clusters restricted to harmonic 5 occurred during the early and middle portions of the stimulus. Overall, across trajectories involving changes in formant frequency, the harmonic location of significant PLV_TFS_ in yNH generally shifted in the same direction as the formant, such that increases in formant frequency were accompanied by a shift toward higher harmonics and decreases by the complementary shift toward lower harmonics. In the harmonic marginals, PLV_TFS_ was significantly above noise at harmonics 5-6 in yNH for all trajectories except *BWinc*, for which significant above-noise PLV_TFS_ was restricted to harmonic 5; in mNH, significant clusters spanned harmonics 5-6 for *FFdec*, *FFinc*, and *FFBWdec*, but were restricted to harmonic 5 for *BWdec*, *BWinc*, and *FFBWinc* (all cluster-corrected *p*< 0.05). In contrast, no significant above-noise temporal marginal clusters were found for any trajectory in either group.

In the harmonic marginals, PLV_TFS_ was significantly higher in yNH than in mNH at harmonic 3 for *FFBWinc* (*p*= 0.029; **Fig. 2C**). No significant between-group differences were observed for the remaining trajectories. In the temporal marginals, no significant between-group clusters were found for any of the six formant trajectories.

To further quantify the age differences observed in PLV, subsequent analyses focused on summary metrics derived from the harmonicgrams. These included mean PLV_ENV_ and mean PLV_TFS_, together with dynamic measures capturing changes in phase locking over the course of the stimulus: the slope of the temporal marginal for PLV_ENV_ and the slope of the PLV_TFS_ spectral centroid. The following sections examine age-related differences in these derived metrics.

### 3.2 Mean PLV_ENV_, but not Mean PLV_TFS_, decreased with age

Previous FFR studies have often summarized responses using scalar measures combining information across harmonics (e.g., Vasilkov et al., 2021; Ponsot et al., 2025). Although our stimuli varied dynamically, we could similarly obtain a single measure of overall phase locking by averaging PLV across all harmonics and time bins of each harmonicgram (see ***Section 2.2.5***).

Mean PLV_ENV_ was negatively associated with age across all eight stimulus trajectories, and all associations remained significant after FDR correction (**Fig. 3A**; all *p*< 0.05).

**Figure 3.**
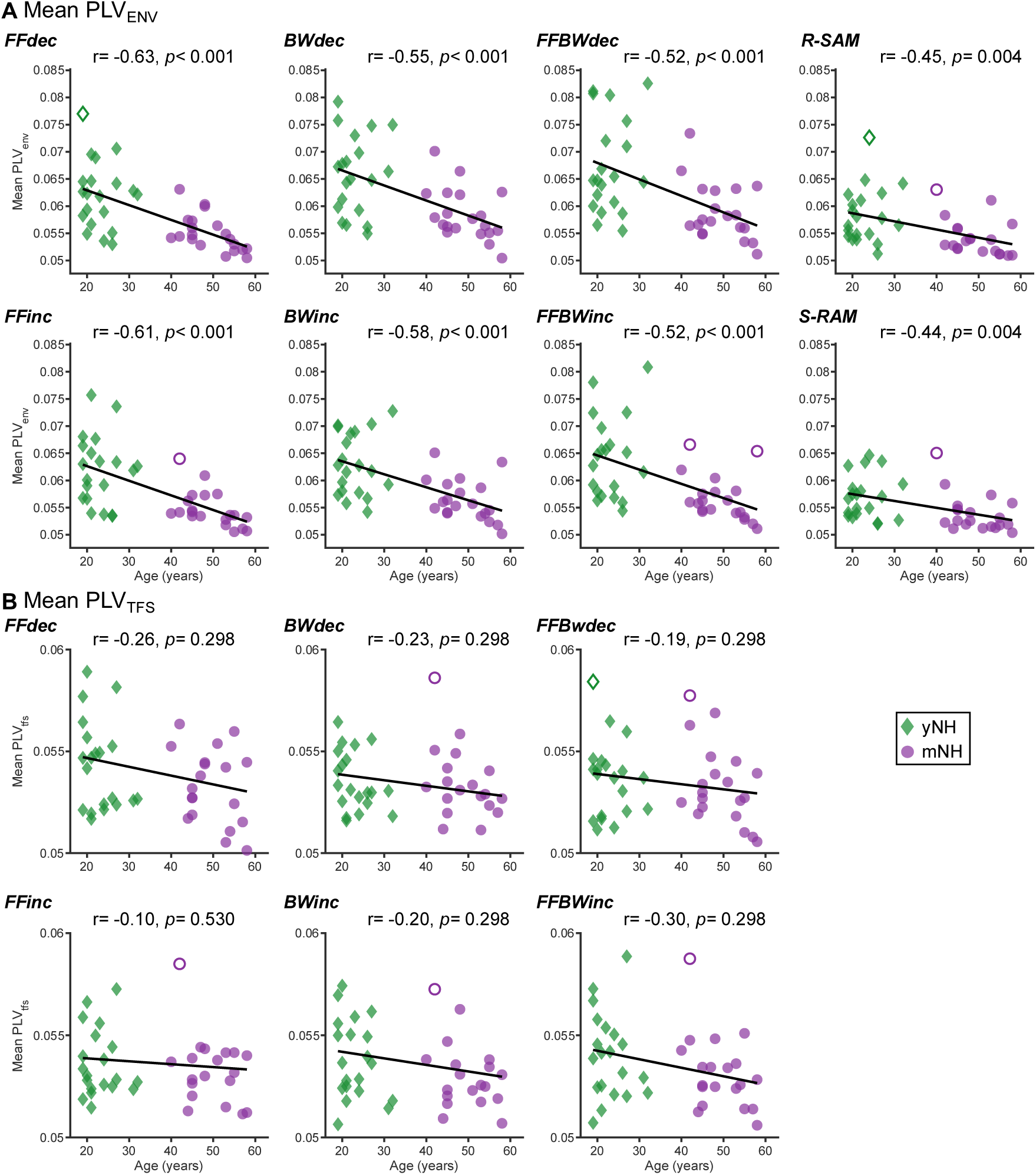
Associations between age and mean PLV. ***A***, Individual mean envelope-related phase-locking values (PLV_ENV_) as a function of age for the eight stimulus trajectories. ***B***, Individual mean temporal-fine-structure-related phase-locking values (PLV_TFS_) as a function of age for the six formant-frequency (FF), formant-bandwidth (BW), and combined FF and BW trajectories. Green diamonds represent young normal-hearing (yNH) adults and purple circles represent middle-aged normal-hearing (mNH) adults. Black lines indicate linear fits across all participants. Open symbols denote observations identified by the within-group outlier criterion; these observations were retained in all analyses. Pearson correlation coefficients (r) and corresponding *p* values are reported in each panel; displayed *p* values were adjusted for multiple comparisons using the false discovery rate (FDR).

In contrast, Mean PLV_TFS_ was not significantly associated with age for any of the six formant trajectories after FDR correction (**Fig. 3B**; all *p*≥ 0.298).

### 3.3 Age-related reductions in PLV_ENV_ tracking of formant bandwidth and temporal-envelope shape

Beyond mean PLV, our dynamic stimuli allowed us to examine how PLV_ENV_ changed as formant frequency, formant bandwidth, and temporal-envelope shape varied over the course of the stimulus. The temporal marginal of PLV_ENV_ represents mean phase locking across harmonics at each time point (**Fig. 2B**). For trajectories involving changes in formant bandwidth or temporal-envelope shape, this measure varies systematically with the acoustic feature being manipulated. PLV_ENV_ is higher when the formant bandwidth is broader and when the temporal modulation is more rectangular, because these stimulus configurations produce stronger phase locking at higher harmonics. As the formant bandwidth narrows or the modulation becomes more sinusoidal, phase locking at these higher harmonics decreases, reducing the PLV_ENV_ temporal marginal.

The slope of the temporal marginal therefore provides a measure of how strongly PLV_ENV_ changes as these acoustic features evolve over time in our stimuli. Importantly, differences between age groups in the absolute level of the temporal marginal do not necessarily imply differences in this dynamic measure. This distinction is illustrated in the PLV for the FF trajectories: changes in formant frequency in the absence of changes in formant bandwidth produced little systematic variation in the PLV_ENV_ temporal marginal. Because formant bandwidth remained constant, there was no corresponding increase or decrease in the contribution of higher-order harmonics to the mean PLV_ENV_ across harmonics, resulting in temporal slopes close to zero in both groups even though the two age groups differ in their mean PLV_ENV_ across the temporal marginal (see *FFdec* and *FFinc* panels in **Fig. 2B**)

Consistent with this interpretation, mean slopes for the FF trajectories remained close to zero and were not significantly associated with age (all *p*≥ 0.071; **Fig. 4**). In contrast, mean slope was significantly associated with age for *BWdec*, *FFBWdec*, *R-SAM*, and *S-RAM* (all *p*≤ 0.026). In each case, increasing age was associated with slopes closer to zero: negative slopes became less negative for the decreasing trajectories and *R-SAM*, whereas the positive slope for *S-RAM* became smaller. Thus, increasing age was associated with a weaker change in PLV_ENV_ as formant bandwidth or temporal-envelope shape evolved over time, consistent with reduced subcortical neural tracking of these dynamic acoustic changes. Associations with age were not significant for *BWinc* and *FFBWinc* (all *p*≥ 0.069).

**Figure 4.**
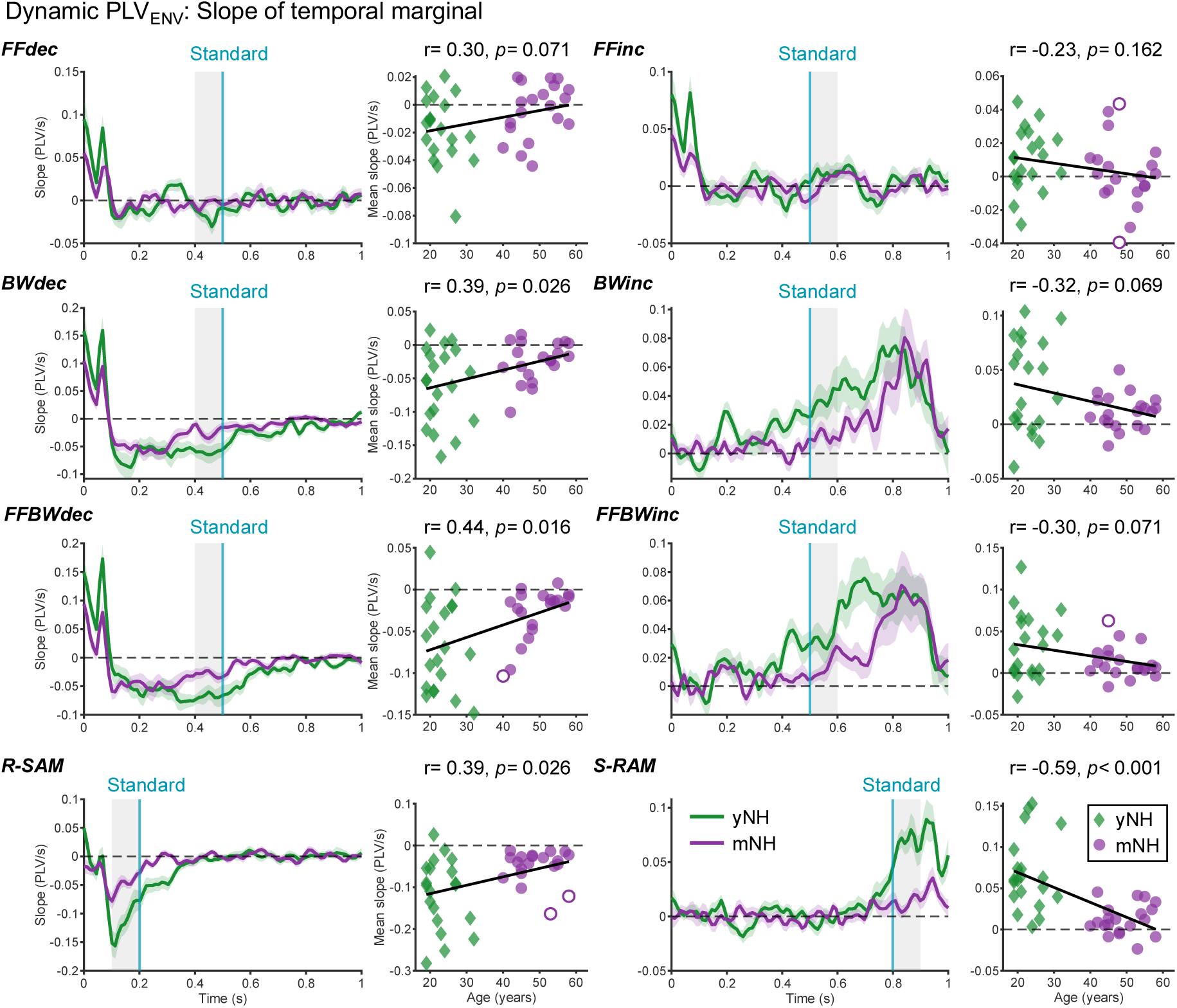
Age-related differences in the temporal dynamics of PLV_ENV_. For each of the eight stimulus trajectories, the temporal derivative of the PLV_ENV_ temporal marginal is shown together with the association between the corresponding individual mean slope and age. Green and purple curves indicate group-mean slopes for young normal-hearing (yNH) and middle-aged normal-hearing (mNH) adults, respectively; shaded bands indicate ±1 SEM. Turquoise vertical lines indicate the time at which each dynamic stimulus reached the acoustic value of the standard stimulus used in the corresponding psychoacoustic task. Gray regions indicate the 100-ms analysis windows positioned immediately before or after this time point, depending on trajectory direction, such that the change in the relevant acoustic dimension within the window matched the direction in which the target stimulus approached the standard stimulus during adaptive tracking in the psychoacoustic tasks. Scatter plots show individual mean slopes as a function of age. Green diamonds represent yNH participants and purple circles represent mNH participants. Open symbols denote observations identified as outliers using Tukey’s 1.5 × interquartile range (IQR) criterion, applied separately within each age group and stimulus trajectory; these observations were retained in all analyses. Black lines indicate linear fits across all participants, and horizontal dashed lines indicate zero slope. Pearson correlation coefficients (r) and FDR-adjusted *p* values are reported above each scatter plot.

### 3.4 No age-related differences in PLV_TFS_ tracking of formant frequency and bandwidth

As with PLV_ENV_, our dynamic stimuli allowed us to examine how PLV_TFS_ changed as formant frequency and bandwidth varied over time. For PLV_TFS_, we assessed this tracking through changes in the distribution of PLV_TFS_ across harmonics, quantified by the slope of the PLV_TFS_ spectral centroid (see ***Section 2.2.5***).

The PLV_TFS_ harmonicgrams showed that significant phase locking was concentrated around the fifth harmonic, corresponding to the 565-Hz formant frequency (**Fig. 2C**). The PLV-weighted spectral centroid therefore summarizes the distribution of PLV_TFS_ across harmonics at each time point. When formant frequency changes, the centroid shifts across harmonics in the same direction as the formant, allowing its temporal evolution to capture subcortical neural tracking of the dynamic formant-frequency trajectory. The centroid is also sensitive to changes in formant bandwidth: broader bandwidths increase the contribution of higher-order harmonics to PLV_TFS_, shifting the centroid toward higher frequencies, whereas narrower bandwidths reduce this contribution. Thus, the slope of the PLV_TFS_ spectral centroid across the stimulus interval provides a measure of the dynamic redistribution of PLV_TFS_ across harmonics during changes in formant frequency and bandwidth, allowing us to test whether this tracking varies with age.

Consistent with this reasoning, group-mean PLV_TFS_ spectral centroids decreased over time for all three decreasing trajectories and increased for all three increasing trajectories (**Fig. 5**). Linear fits captured these temporal changes reasonably well across conditions and groups (R² = 0.57-0.97).

**Figure 5.**
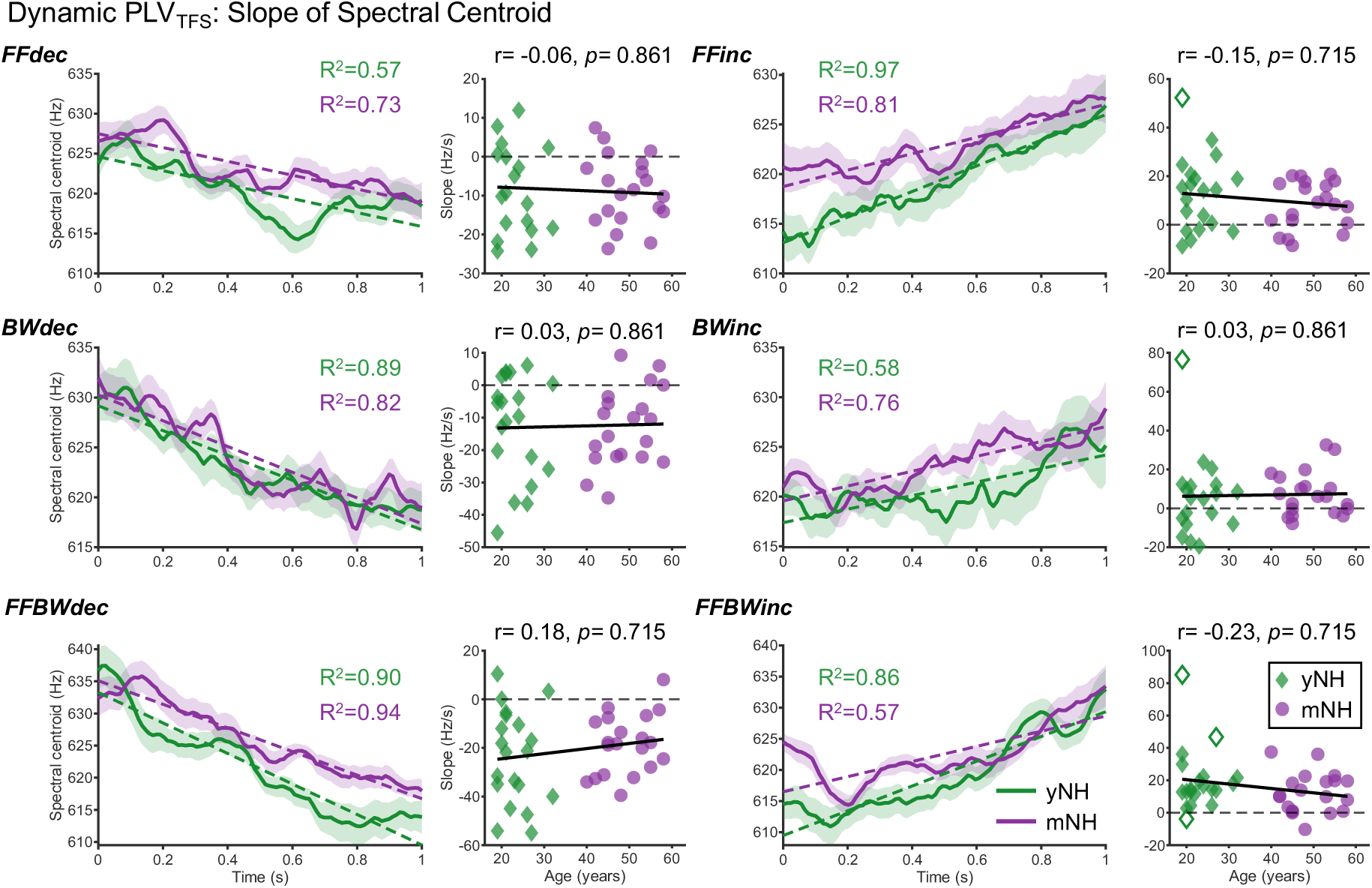
Age-related differences in the temporal dynamics of PLV_TFS_. For each of the six formant trajectories, the temporal evolution of the PLV_TFS_ spectral centroid is shown together with the association between the corresponding individual centroid slope and age. Green and purple curves indicate group-mean spectral centroids for young normal-hearing (yNH) and middle-aged normal-hearing (mNH) adults, respectively; shaded bands indicate ±1 SEM. Dashed colored lines indicate linear fits to the group-mean spectral-centroid trajectories, with the corresponding coefficients of determination (R²) shown in matching colors. Scatter plots show individual spectral-centroid slopes as a function of age. Green diamonds represent yNH participants and purple circles represent mNH participants. Open symbols denote observations identified as outliers using Tukey’s 1.5 × interquartile range (IQR) criterion, applied separately within each age group and stimulus trajectory; these observations were retained in all analyses. Participant-trajectory fits were excluded due to poor fitting (see **Section 2.2.5**). Black lines indicate linear fits across all participants, and horizontal dashed lines indicate zero slope. Pearson correlation coefficients (r) and FDR-adjusted *p* values are reported above each scatter plot.

Despite this clear temporal tracking, individual spectral-centroid slopes were not significantly associated with age for any of the six trajectories after FDR correction (all *p*≥ 0.715). Thus, although PLV_TFS_ spectral centroids tracked the dynamic changes in formant frequency and bandwidth, the rate of this redistribution across harmonics showed no evidence of an association with age.

### 3.5 No associations between age and psychoacoustic discrimination performance

The age-related changes in PLV measures described above raise the question of whether perceptual sensitivity to the corresponding acoustic features also changes with age. To address this, we measured discrimination thresholds for formant frequency, formant bandwidth, and temporal-envelope shape. Thresholds were assessed in quiet and in background noise to examine whether associations with age became more apparent under more challenging listening conditions.

In quiet, discrimination thresholds were not significantly associated with age for any of the three tasks (all *p*≥ 0.721; **Fig. 6A**). The small correlation coefficients between thresholds in quiet and age could be partly explained by the limited variability of these thresholds, which were concentrated at relatively low values, particularly for FF and BW discrimination.

**Figure 6.**
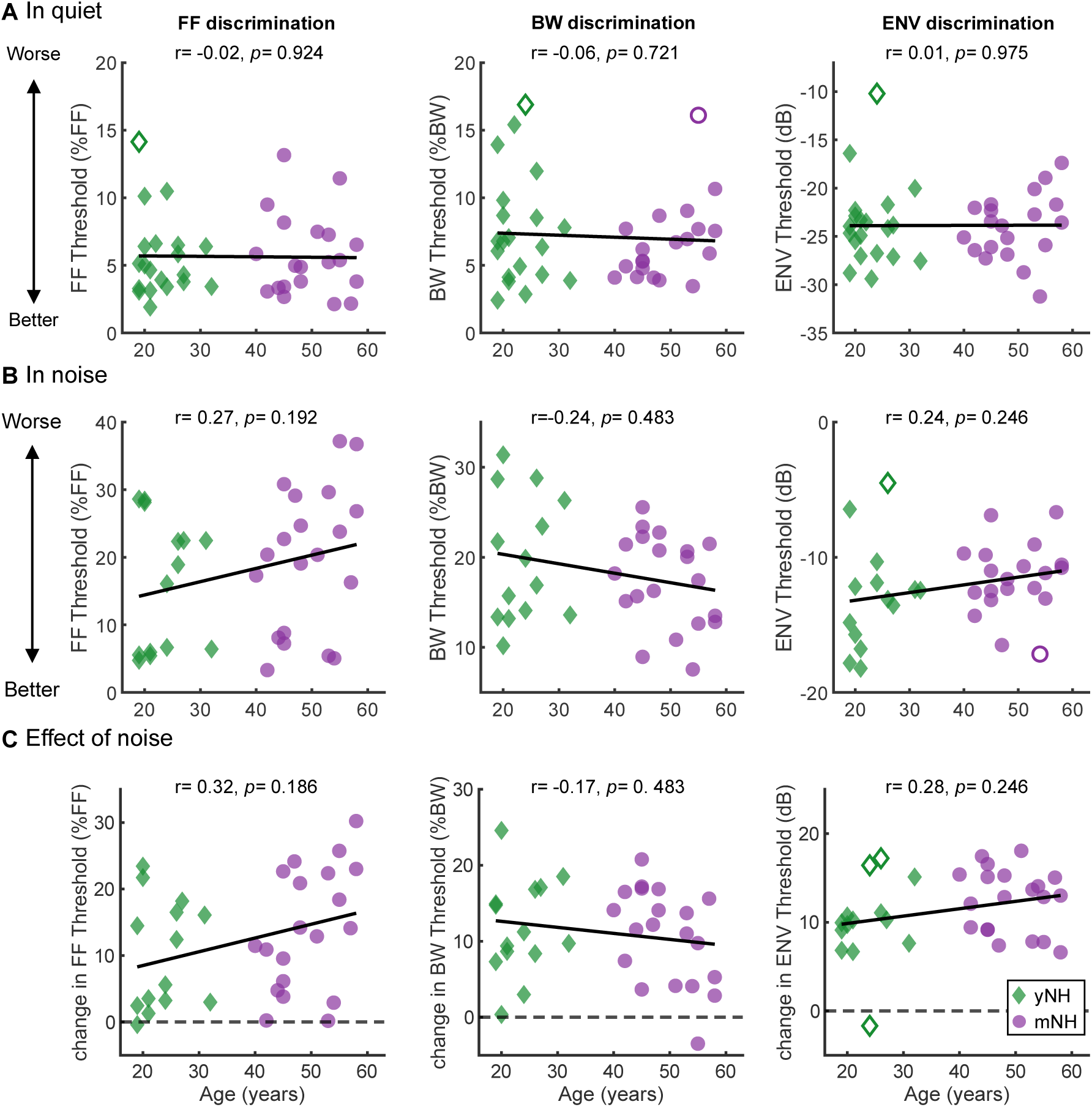
Associations between age and psychoacoustic discrimination thresholds in quiet and noise. ***A***, Discrimination thresholds measured in quiet. ***B***, Discrimination thresholds measured in noise. **C**, Effect of noise, expressed as the difference between thresholds measured in noise and quiet (noise -quiet). Columns correspond to formant-frequency (FF), formant-bandwidth (BW), and envelope-shape (ENV) discrimination. Lower thresholds indicate better discrimination performance. In *C*, horizontal dashed lines indicate no change in threshold between noise and quiet; positive values indicate poorer performance in noise. Green diamonds represent young normal-hearing (yNH) participants and purple circles represent middle-aged normal-hearing (mNH) participants. Open symbols denote observations identified as outliers using Tukey’s 1.5 × interquartile range (IQR) criterion, applied separately within each age group and task-condition combination; these observations were retained in all analyses. Black lines indicate linear fits across participants. Pearson correlation coefficients (r) and FDR-adjusted *p* values are reported above each scatter plot. False discovery rate correction was performed separately for the FF, BW, and ENV discrimination tasks using the Benjamini-Hochberg procedure.

In noise, FF and BW thresholds showed greater variability, and correlations with age were stronger in absolute magnitude across all three tasks, although none reached statistical significance (all *p*≥ 0.192; **Fig. 6B**).

No significant associations with age were observed for the effect of noise (noise-quiet threshold) in any of the three tasks (**Fig. 6C**). For FF and envelope-shape discrimination, the effect of noise tended to become larger with increasing age (FF: r= 0.32, *p*= 0.186; envelope shape: r= 0.28, *p*= 0.246), although neither association reached statistical significance. BW discrimination showed the opposite descriptive pattern, with the effect of noise tending to decrease with age (r= -0.17, *p*= 0.483), but this association was also not significant.

### 3.6 PLV metrics did not significantly predict psychoacoustic discrimination thresholds

To investigate the relationship between PLV metrics and perceptual performance, we examined whether individual differences in FFR metrics were associated with discrimination thresholds for formant frequency, formant bandwidth, and temporal-envelope shape. Importantly, our dynamic stimuli allowed us to assess not only mean PLV but also how PLV tracked changes in the same acoustic features evaluated psychophysically. We examined these associations after accounting for listening condition, both before and after adjustment for age.

Listening condition (Quiet vs Noise) accounted for a substantial proportion of the variance in psychoacoustic performance, with marginal R² values of 43.14%, 58.34%, and 73.09% for FF, BW, and ENV discrimination, respectively (**Fig. 7**). Relative to the intercept-only models, adding condition improved fit for FF (χ²(1)= 44.77), BW (χ²(1)= 65.23), and ENV (χ²(1)= 101.19; all p< 0.001). Adding age increased marginal R² by 1.77, 0.99, and 0.40%, respectively, but did not significantly improve model fit for any task (FF, χ²(1)= 1.61, *p*= 0.204; BW, χ²(1)= 1.63, *p*= 0.202; ENV, χ²(1)= 0.77, *p*= 0.381).

**Figure 7.**
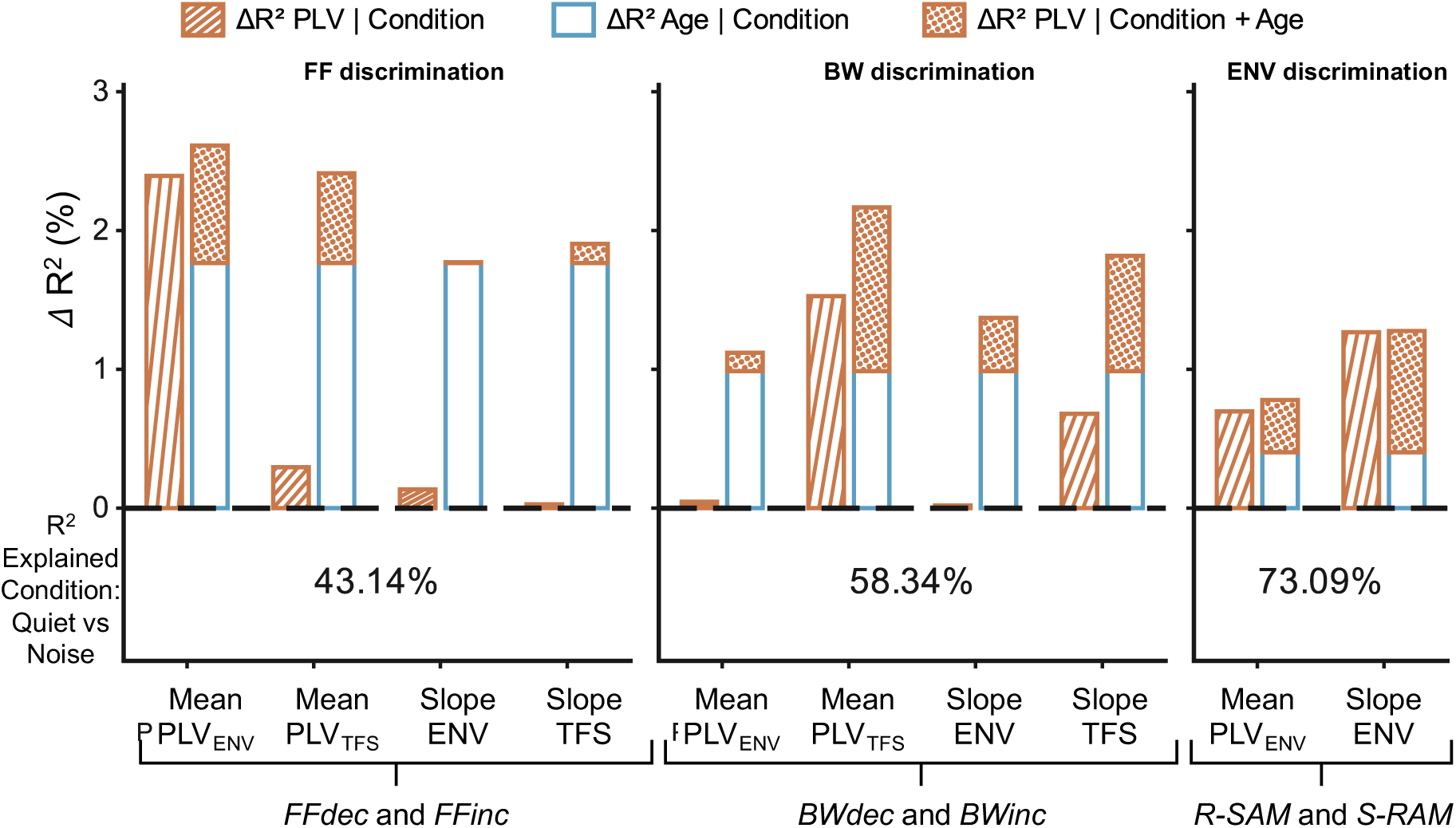
Incremental variance explained by PLV metrics and age in psychoacoustic discrimination thresholds. Incremental marginal R² (ΔR²) values are shown for formant-frequency (FF), formant-bandwidth (BW), and envelope-shape (ENV) discrimination. Striped, orange bars indicate the additional variance explained by each PLV predictor after accounting for listening condition (quiet vs noise). Blue bars indicate the additional variance explained by age after accounting for condition, and dotted orange segments indicate the further variance explained by the PLV predictor after accounting for both condition and age. For FF and BW discrimination, PLV predictors included mean PLV_ENV_, mean PLV_TFS_, PLV_ENV_ temporal slope, and PLV_TFS_ spectral-centroid slope; for ENV discrimination, predictors included mean PLV_ENV_ and PLV_ENV_ temporal slope. The marginal R² of the condition-only model is reported below each task.

After accounting for listening condition, the additional variance explained by individual EEG predictors ranged from 0.02 to 2.39% across tasks (**Fig. 7**). The largest increment was observed for mean PLV_ENV_ in FF discrimination (ΔR²= 2.39%, χ²(1)= 2.27, *p*= 0.528), followed by mean PLV_TFS_ in BW discrimination (ΔR²= 1.53%, χ²(1)= 2.42, *p*= 0.478). None of the PLV predictors explained significant additional variance after FDR correction.

After accounting for both listening condition and age, PLV-related increments in marginal R² ranged from 0.01 to 1.18% and remained nonsignificant. The largest age-adjusted increment was observed for mean PLV_TFS_ in BW discrimination (ΔR²= 1.18%, χ²(1)= 1.88, *p*= 0.490). For mean PLV_ENV_ in FF discrimination, the additional variance explained by the PLV predictor was 0.85% after adjustment for age (χ²(1)= 0.86%, *p*= 0.779), compared with 2.39% when only listening condition was included in the model. Similarly, for the PLV_ENV_ temporal slope in ENV discrimination, the age-adjusted PLV increment was 0.87% (χ²(1)= 2.02, *p*= 0.311), compared with 1.27% before adjustment for age. Several other PLV predictors also showed smaller increments after age was included in the model. This reduction in incremental variance after adjustment for age is consistent with the possibility that part of the variance in psychoacoustic performance associated with the PLV metrics overlaps with variance explained by age, limiting their additional explanatory value once age is included in the model.

Overall, the PLV metrics did not explain significant additional variance in psychoacoustic discrimination thresholds beyond that accounted for by listening condition, either before or after adjustment for age.

## 4 Discussion

We combined FFR recordings to dynamic vowel-like and amplitude-modulated sounds with psychophysical measures along the same acoustic dimensions in yNH and mNH adults. In mNH, PLV_ENV_ was reduced for both low-and high-frequency stimuli, with lower values across harmonics and weaker tracking of changes in formant bandwidth and envelope shape (**Figs. 2-4**). In contrast, we found no significant age-related difference in PLV_TFS_, even though it tracked changes in formant frequency and bandwidth in both groups (**Figs. 2C, 3B, 5**). Discrimination thresholds were not significantly associated with age, in quiet or in noise, and FFR measures did not significantly explain individual differences in performance (**Figs. 6, 7**). These findings indicate that age-related changes in subcortical envelope coding can occur without corresponding changes in perceptual sensitivity.

### 4.1 Reduced envelope coding across stimuli and cochlear regions in middle age

The reduction in PLV_ENV_ for both low-frequency single-formant complexes and high-frequency amplitude-modulated tones is consistent with previous reports of weaker envelope coding with age (Vasilkov et al., 2021; Märcher-Rørsted et al., 2022; Garrett et al., 2025), including in middle-aged adults (Ponsot et al., 2025; Zink et al., 2025). Here, we extend these findings to dynamic stimuli designed to target apical and basal cochlear regions.

At high frequencies, age-related differences in PLV_ENV_ were concentrated in the portions of the amplitude-modulated stimuli with more rectangular, rather than sinusoidal, envelopes (**Fig. 2B**). This is consistent with stronger age effects reported for the high-frequency components of the envelope (Vasilkov et al., 2021; Garrett et al., 2025; Zink et al., 2025).

At low frequencies, age-related differences in PLV_ENV_ were mostly restricted to the lowest harmonics when only formant frequency changed, but extended to higher harmonics when formant bandwidth was broader (**Fig. 2B**). The greater contribution of higher harmonics previously observed with rectangular amplitude modulation (Vasilkov et al., 2021; Garrett et al., 2025; Ponsot et al., 2025) was thus paralleled by greater PLVs extending to higher harmonics for vowel-like sounds with broader bandwidth. These results refine our previous findings (Ponsot et al., 2025): age-related reductions in envelope coding were present for all trajectories and were most extensive for high-frequency rectangular amplitude-modulated tones and for low-frequency vowel-like sounds with rich spectra (**Figs. 2B, 3A**). This result is consistent with histological evidence that age-related AN fiber loss is distributed across the cochlea (Wu et al., 2019). Vowel-like sounds with rich spectra could therefore be explored as FFR-based markers of CS at low frequencies (Ponsot et al., 2025), complementing the high-frequency RAM stimuli used so far (Vasilkov et al., 2021; Van Der Biest et al., 2023; Garrett et al., 2025). However, the relative contributions of different cochlear regions to FFR_ENV_ cannot necessarily be inferred from the spectral content of the stimulus alone (Zhu et al., 2013; Keshishzadeh et al., 2020) and further studies are therefore needed to establish the cochlear frequency specificity of broad-bandwidth formant stimuli and RAM-evoked FFR_ENV_.

Beyond mean PLV_ENV_, the dynamic stimuli allowed us to examine how PLV_ENV_ tracked changes in formant frequency, formant bandwidth and envelope shape, and whether this tracking varied with age (**Fig. 4**). When formant bandwidth narrowed, alone or together with formant frequency, or when envelope shape changed between rectangular and sinusoidal modulation, the slope of the PLV_ENV_ temporal marginal was closer to zero in older participants, indicating weaker tracking in mNH adults. Mean PLV_ENV_ and slope thus carry complementary information: for the formant-frequency trajectories, mean PLV_ENV_ decreased with age (**Fig. 3A**), whereas the slope did not (**Fig. 4**). This highlights the importance of measuring how FFRs track changes in acoustic features, as such measures provide information complementary to conventional overall FFR metrics and are also sensitive to age-related changes.

### 4.2 Temporal fine structure coding is largely preserved in middle age

In contrast, we found no significant age-related difference in PLV_TFS_. PLV_TFS_ was concentrated around harmonics 5-6, near the formant peak, and PLV_TFS_ spectral centroid tracked changes in formant frequency and bandwidth in both age groups (**Figs. 2C, 5**). Neither mean PLV_TFS_ nor the PLV_TFS_ centroid slope was significantly associated with age (**Figs. 3B, 5**). These findings suggest that TFS coding is less affected than envelope coding in middle-aged adults, consistent with intact AN phase locking to fine structure in aged gerbils (Steenken et al., 2026). However, these findings contrast with the age-related FFR_TFS_ reductions reported for low-frequency tones (Märcher-Rørsted et al., 2022, 2026) and for vowel-like stimuli (Anderson et al., 2012; Ponsot et al., 2025).

Age may partly explain this discrepancy: some of these cohorts were older (48-76 years in Märcher-Rørsted et al., 2022; up to 77 years in Märcher-Rørsted et al., 2026), the proportion of AN fibers lost increases with age (Wu et al., 2019), and effects on FFR_TFS_ are expected to be small in middle age (Märcher-Rørsted et al., 2022). Age cannot, however, explain the discrepancy with our previous study, whose older group (47 ± 6 years) was comparable to the present mNH group (Ponsot et al., 2025). That study used stationary rather than dynamic stimuli, a higher sound level (75 vs. 62.5 dB SPL), and about four times more repetitions per participant (about 2850 vs. 840), which may have favored the detection of small effects.

In sum, the ENV and TFS components of the FFR thus appear to index different aspects of neural coding. ENV-based markers may be particularly sensitive to early aging (and hypothesized CS) because they reflect the envelope beating produced by harmonics across a wide range of cochlear channels (Zhu et al., 2013).

### 4.3 No age effect on discrimination thresholds

Discrimination thresholds for formant frequency, formant bandwidth and envelope shape were not significantly associated with age, in quiet or in background noise (**Fig. 6**). Background noise increased thresholds in all three tasks, but its effect was not significantly associated with age (**Fig. 6C**). For formant frequency, this result agrees with reports that discrimination near F1, unlike near F2, is robust to aging (Coughlin et al., 1998) and to hearing loss (Carney et al., 2023; Mishra et al., 2023). This result is also in line with the largely negative literature searching for perceptual correlates of age-related synaptopathy (Prendergast et al., 2019; Patro et al., 2021).

Detecting perceptual consequences of CS is believed to require tasks that (i) place substantial demands on peripheral coding under challenging listening conditions, while minimizing cognitive contributions (DiNino et al., 2022), and (ii) preferentially recruit low-and medium-spontaneous-rate (L/MSR) AN fibers (Bharadwaj et al., 2019). Our study may have fallen short on both counts. First, although inter-individual variability was larger in noise than in quiet, it remained limited, and the smaller sample in noise (n=34) may have prevented the detection of subtle effects (Bramhall and McMillan, 2024). Second, our tasks may not have shifted coding sufficiently toward L/MSR fibers (Bharadwaj et al., 2015), even at the higher level and in noise, and the contribution of the different AN fiber types to perception is still debated (Carney, 2018). The absence of significant associations therefore does not rule out perceptual consequences of CS: it suggests that, if present, they are subtle in middle age, which makes a sensitive behavioral test usable in the clinic difficult to envisage. An additional possibility is that discrimination of spectral and temporal features in complex sounds at conversational speech levels is largely unaffected by aging or CS. If CS-related perceptual deficits become apparent only at higher presentation levels, their ecological relevance may be limited.

Perceptual consequences of CS in middle age may also appear first as increased listening effort rather than lower accuracy (Zink et al., 2025; Zhen et al., 2025), and further work based on specifically tailored measurements, with pupillometry or reaction time, is needed.

### 4.4 No significant relationship between FFR and psychophysical performance

Despite deriving FFR metrics sensitive to changes in the same features assessed psychophysically (formant frequency, formant bandwidth and envelope shape), we found no significant association between neural responses and discrimination thresholds (**Fig. 7**). Previous studies relating subcortical FFRs to perception have yielded mixed results. No association between ENV-or TFS-based FFR metrics and perceptual measures has been reported for temporal-envelope processing (Patro et al., 2021), spectral-envelope processing (Ponsot et al., 2025), speech-in-noise perception (Schoof and Rosen, 2016) or vowel recognition (Berg et al., 2025). Others found significant relationships with temporal-envelope processing (Bharadwaj et al., 2015), vowel identification (Won et al., 2016) or speech-in-noise performance (Mai et al., 2018; Mepani et al., 2021; Garrett et al., 2025; Zink et al., 2025). Although matching stimuli across FFR and psychoacoustic measurements has been proposed as a way to clarify these discrepancies (Garrett et al., 2025), our results suggest that assessing the same acoustic features does not necessarily reveal an association between subcortical neural responses and perceptual performance.

One possible explanation for the lack of correlation between psychophysical and FFR responses is that central gain could compensate for reduced peripheral activity by enhancing responses at higher stages of the auditory system, thereby helping to preserve perceptual sensitivity. Märcher-Rørsted et al. (2026) reported age-related reductions in ABR wave I and FFRs, alongside preserved ABR wave V and enhanced cortical responses, suggesting increased brainstem gain for transient responses. However, these cortical changes were not significantly correlated with the cochlear measures, and they may reflect brain aging, including reduced inhibition, rather than compensation for peripheral loss. Future studies combining subcortical and cortical measures, such as the acoustic change complex, which is associated with psychophysical discrimination thresholds (Meehan et al., 2025; Rekswinkel et al., 2026), could help determine whether central gain contributes to preserving perception in middle age.

Appendix: Noise-floor estimation for phase-locking values

Phase-locking values can remain above zero even when responses are not phase-locked to the stimulus; their expected values also depend on the number of available epochs. Simulated null distributions were therefore used to assess whether the observed PLV responses exceeded values expected under the noise model.

Null distributions were generated separately for each age group and stimulus trajectory using Gaussian white-noise epochs. Simulated recordings comprised 1-s epochs sampled at 16,384 Hz, with epoch-pool sizes matched to the corresponding participant and trajectory. The retained counts ranged from 382 to 812 epochs. Each pool was divided equally between polarities, with odd totals reduced by one. Phases were extracted at the exact frequencies of harmonics 1-10 of the 113-Hz fundamental using 100-ms Blackman-Harris windows with 90% overlap. Each simulated participant’s PLVs were estimated from 100 bootstrap samples of 400 epochs, with equal numbers drawn from each polarity pool. These participant-level estimates were then averaged to obtain one group-level null realization. This procedure was repeated to generate 200 independent realizations for each group and trajectory. The same null distributions were used for PLV_ENV_ and PLV_TFS_ because a fixed polarity-related phase shift does not change a uniform circular phase distribution.

Each null realization consisted of a harmonic-by-time map of the group-mean PLV expected under the simulated noise model. Harmonic and temporal marginal curves were derived from these maps using the same averaging operations as for the observed data. Consequently, each observed group map or marginal was accompanied by 200 null maps or curves of the same dimensions.

## Ethical Approvals and Consent to Participate

All experiments were approved by the Institut Europeen d’Administration des Affaires (INSEAD). The study was conducted in accordance with the Declaration of Helsinki. All subjects were informed about the experimental procedures, provided written informed consent, and received financial compensation for their participation.

## Acknowledgments

This work was funded by the Agence Nationale de la Recherche (Grant No. ANR-22-CE28-0010) and the CNRS MITI (EP) and by NIH-NIDCD-R01-DC010813 (LHC).

## CRediT Author Contributions

DLR: Conceptualization, Methodology, Investigation, Formal analysis, Writing -original draft, Writing -review & editing. VB: Conceptualization, Methodology, Investigation, Formal analysis. EP: Conceptualization, Methodology, Formal analysis, Resources, Supervision, Writing -review & editing, Funding acquisition. LHC: Methodology, Formal analysis, Writing -review & editing.

## Data and Code Availability

Data supporting the findings of this study are available from the corresponding author upon reasonable request. Codes used for analyses reported in this paper are available from the authors upon reasonable request.

